# Serotypic heterogeneity in the response to pneumococcal vaccine

**DOI:** 10.64898/2026.02.06.704301

**Authors:** Gatien Durand, Camillia Belhoul, Merieme Bensalah, Maxime Jeljeli, Laurie Toullec, Marine Gil, Elodie Lachiche, Camille Baron, Claire Goulvestre, Christian Drouet, Frédéric Batteux, Lucie Chevrier

**Affiliations:** Université Paris Cité, Faculté de Médecine, AP-HP-Centre Université de Paris, Hôpital Cochin, Service d’immunologie biologique, 75679 Paris, France; Département 3I « Infection, Immunité et Inflammation », Institut Cochin, INSERM U1016, Université Paris Cité, 75679 Paris, France; Université Grenoble Alpes, 38400 Saint-Martin-d’Hères, France; Institut Pasteur de Lille, Lille, France

**Keywords:** Pneuomoccal vaccine, ELISA, OPA, hetereogeneity, serotypes

## Abstract

The assessment of pneumococcal vaccine response currently relies on a single IgG concentration threshold, identical for all serotypes, as recommended by the WHO and AAAAI. However, the recommended thresholds do not take the wide inter-serotype variability into account. The purpose of this study was to determine if the antibody threshold linked to a functional immune response varies according to serotype.

We performed a retrospective analysis on 729 samples from adults at risk of invasive pneumococcal disease (IPD), sent between 2018 and 2024 to assess vaccine response. Specific IgG concentrations for seven vaccine serotypes were measured by ELISA and compared to opsonophagocytic assay (OPA) results, a functional test considered as the gold standard. For each serotype, we determined the most predictive IgG concentration (PIC) for a positive OPA result using ROC curves, Youden’s index, and bootstrap analysis with 1,000 resamples. The resulting PIC were then compared using non-parametric tests (Kruskal-Wallis test followed by Dunn’s post-hoc test with Holm’s correction).

The PIC varied considerably among serotypes, ranging from 0.84 to 4.74 µg/mL. This variability was found to be statistically significant (p<0.0001). Areas Under the Curve (AUC), ranging from 0.73 to 0.87, demonstrate good diagnostic performance. Overall, the application of serotype-specific thresholds in patients significantly change the classification of vaccination status compared to a single threshold (Cochran, McNemar).

These results indicate that the protective antibody threshold is not universal. A serotype-specific approach would allow a more precise and relevant assessment of the pneumococcal vaccine response.

**Author summary:** The introduction of the 7-valent pneumococcal conjugate vaccine (PCV-7) in the early 2000s significantly changed the epidemiology of invasive pneumococcal disease by modifying serotype distribution. However, vaccine-induced immunity varies across serotypes, and this heterogeneity remains incompletely understood. In this study, we first assessed differences in vaccine responses according to serotype. We then determined predictive serotype-specific immunoglobulin concentration for the seven routinely serotypes tested in our laboratory, defined as the minimal levels required to induce a positive opsonophagocytic activity (OPA) response. The results enable a more accurate assessment of serotype-specific vaccine immunity, supporting improved patient stratification and guiding booster vaccination in individuals with insufficient responses.

## Introduction

*Streptococcus pneumoniae* (commonly named pneumococcus) is a major cause of infections producing different clinical manifestations(1). Local spread, aspiration or blood dissemination of pneumococcus is responsible for invasive pneumococcal diseases (IPD)(2), with variable clinical picture according to the location of the bacterial dissemination(3). Pneumococcus is both the leading cause of bacterial pneumonia, with exacerbated morbidity and mortality in developing countries in children under five(4) and proportionally the main causative agent of meningitis mortality across all age groups(5). One of the most efficient ways of combating IPD is to develop vaccines targeting polysaccharide capsule(6), which is also highly heterogenous(7) and is the main virulence factor(8). Polysaccharide antigens stimulate populations of B lymphocytes independently of T cells(9), prompting to the development of conjugated vaccines (PCV)(10). The first PCV introduced in the early 2000s targeted the seven most virulent serotypes (4, 6B, 9V, 14, 18C, 19F and 23F)(11). In parallel to the fall of IPD by pneumococcus groups corresponding to vaccine serotypes(11), an increase for non-vaccine serotype prompted to a sequential broadening of the PCV from 7 to 20 valences(12–15). Monitoring the vaccination’s status of patients is crucial, particularly for those who are vulnerable to pneumococcal infections(16). To this aim, two serotype-specific analytical techniques have been validated as gold standard by the World Health Organization (WHO): an enzyme-linked immunosorbent assay (ELISA) to measure antibody titers(17) and an opsonophagocytic assay (OPA) to estimate the functional immunogenicity of specific serotype antibodies(18). The OPA assay monitors the capacity of the patient’s antibodies to develop an effective and specific opsonisation of the bacterial vaccine strain(19). OPA is particularly relevant in monitoring immunocompromised patients in order to decipher his/her efficient immune protection(20). According to WHO and expert consensus, protection cut-off for antibody titer is 1 µg/mL for adult (CAPITA)(21) and 0.35 µg/mL for children(22) for all serotypes. Protection cut-off for OPA corresponds to the lower limit of quantification (LLOQ) determined in every laboratory. Global protection to IPD is achieved when a patient develops a response for at least 70% of the tested serotypes. However, in 2012, the American Academy of Asthma, Allergy and Immunology (AAAAI) proposed a new protection cut-off at 1.3 µg/mL from ELISA for patients with primary immunodeficiency diseases(23). For clinical evaluation of pneumococcal vaccine protection, the protective cut-off for adults is therefore matter of debate. In fact, protective thresholds (1 µg/mL according to WHO or 1.3 µg/mL according to AAAAI) remain empirical and subjective. Moreover, the variability of response between each serotype has not been considered. Seventy nine percent of a healthy non-vaccinated adult cohort displays a median IgG levels equal or greater than 1.3 µg/mL for 74% of polysaccharide specificities tested(24), suggesting the variability of baseline IgG levels. The high variability between serotypes was confirmed by investigating a cohort of 100 healthy controls, where serotype-specific cut-off was determined after vaccination by PPV23(25). Twenty serotypes out of the 23 tested exhibit a cut-off higher than 1 µg/mL, including one serotype with a protective cut-off at 1.3 µg/mL.

The objective of this retrospective study was to investigate serotype heterogeneity in patients at risk of IPD. For each serotype, the specific IgG concentration was measured by ELISA, with values predictive of opsonization capacity according to three approaches: ROC curves, Youden’s index and bootstrap analysis with 1,000 resamples. Our observations were pertaining with a heterogeneity of immune response in a serotype-dependent manner.

## Results

### Cohort description

Between January 2018 and June 2024, 729 samples from adult patients were registered to perform an ELISA and an OPA directed against the 7 serotypes of PCV-7 (4, 6B, 9V, 14, 18C, 19F and 23F). The purpose of these explorations was to determine whether a patient had sufficient vaccine protection at a given time. The median IgG concentration varied according to the serotype, from 0.67 μg/mL for the lowest (serotype 4) to 5.17 μg/mL for the highest (serotype 14) (Table 1).

**Table 1:** Median concentration of vaccine serotype-specific IgG and variability in a series of 729 patient samples.

| Pneumococcus Serotype | Patients (n = 729) |  |
| --- | --- | --- |
|  | ELISA IgG concentration (µg/mL) |  |
|  | Median | 95% CI |
| 4 | 0.67 | 0.13-4.85 |
| 6B | 1.93 | 0.32-18.05 |
| 9V | 1.00 | 0.12-9.10 |
| 14 | 5.17 | 0.54-42.05 |
| 18C | 1.38 | 0.20-11.23 |
| 19F | 2.80 | 0.48-21.85 |
| 23F | 1.30 | 0.17-17.28 |

CI: confidence intervals (P5-P95)

OPA results were displayed in a binary distribution: an OI higher than the LLOQ indicated the production of functional antibodies against the tested serotype. Conversely, an OI lower than the LLOQ highlighted a defective functionality of the antibodies. For patients with a positive OPA, the lowest IgG concentration was 1.06 μg/mL for serotype 4 and the highest IgG concentration 10.24 μg/mL for serotype 14 (Table 2). For patients with a negative OPA, data ranged from 0.46 μg/mL to 2.26 μg/mL, for serotypes 4 et 14 respectively (Table 2). The IgG concentrations between serotypes were then compared by applying a nonparametric Kruskall-Wallis test. We highlighted a heterogeneity of serotype-specific immune response (H(6)=1017, p<0.0001). Due to this high degree of heterogeneity observed between serotypes, we performed a post-hoc Dunn test with Holm correction to identify significant differences between serotypes. Except between serotypes 18C and 23F, where no significant differences were observed, pairwise comparison of the antibody titer heterogeneity was demonstrated between the other serotypes. All data are shown in Figure 1.

**Table 2.** Distribution of anti-polysaccharide IgG concentrations according to OPA test on serum samples.

|  | OPA negative test. | OPA positive test |  |
| --- | --- | --- | --- |
| **Pneumococcus Serotype** | **IgG median concentration (µg/mL) (95% CI)** | **IgG median concentration (µg/mL) (95% CI)** | ***p*-value** |
| **4** | **0.46** (0.11–2.09) | **1.06** (0.20–6.42) | <0.0001 |
| **6B** | **1.32** (0.22–4.47) | **3.12** (0.60–28.48) | <0.0001 |
| **9V** | **0.64** (0.10–3.10) | **1.80** (0.28–12.71) | <0.0001 |
| **14** | **2.26** (0.30–11.15) | **10.24** (2.28–73.71) | <0.0001 |
| **18C** | **0.75** (0.15–3.34) | **2.51** (0.55–15.75) | <0.0001 |
| **19F** | **1.50** (0.28–6.98) | **3.54** (0.66–28.24) | <0.0001 |
| **23F** | **0.73** (0.14–3.83) | **3.71** (0.40–36.67) | <0.0001 |

CI: confidence intervals, Mann-Whitney tests

**Figure 1.**
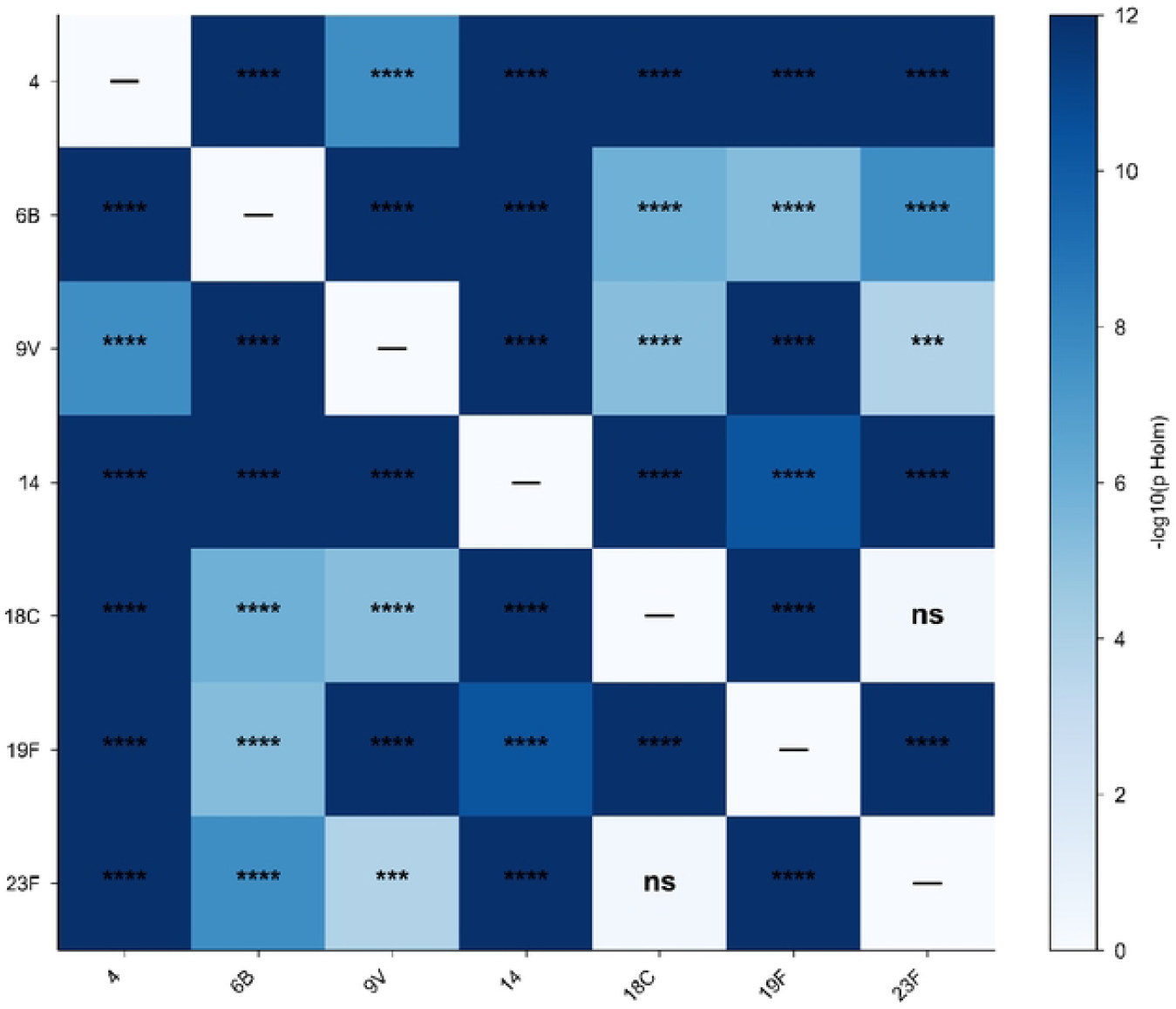
Serotype-dependent heterogeneity of anti-pneumococcal lgG responses. Heatmap showing pairwise differences in median anti-pneumococcal ELISA lgG titers between pneumococcal serotypes. Each cell represents the median titer difference between two serotypes, with color intensity reflecting the magnitude and direction of the effect (scale shown on the right). Overall heterogeneity among serotypes was assessed using a Kruskal-Wallis test followed by Dunn’s multiple comparisons test with Holm correction. Statistical significance is indicated in the matrix (ns, not significant; *** *p*<0.001; **** *p*<0.0001).

### Determining serotype specific predictive IgG concentration

The evidence of heterogeneity prompted us to determine the allocation of IgG concentrations for each serotype. Based on the binary distribution of the OPA results, we next determined for each serotype an IgG concentration that could be predictive of functional antibody (ie, positive OPA result), named PIC for Predictive IgG Concentration.

First, we compared IgG concentration between group with OPA positive and that with OPA negative (Table 2, Figure 2). For all serotypes, a higher titer in ELISA was found for OPA positive results. Due to the observation of a huge range of median values between the different serotypes, we wanted to determine whether there was a significant difference between them. We performed a Kruskal-Wallis test for ELISA titers associated with a negative OPA result (H(6)=496, p<0.0001) and for ELISA titers with a positive OPA result (H(6)=697, p<0.0001), underlying heterogeneity between serotypes in both cases.

**Figure 2.**
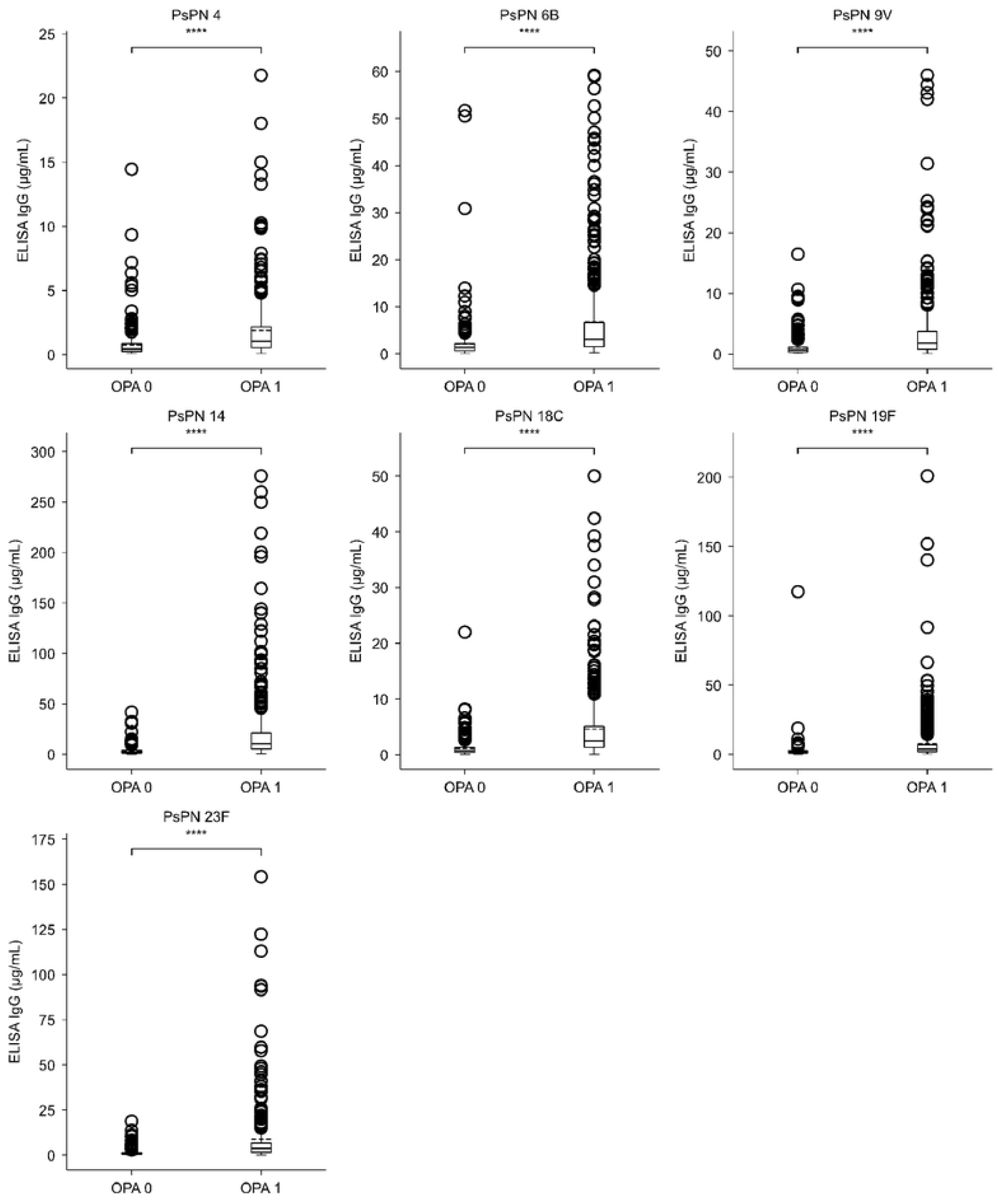
Relationship between ELISA lgG titers and OPA outcome. Anti-pneumococcal ELISA lgG titers were compared between OPA-positive (OPA = 1) and CPA-negative (OPA = O) samples for each serotype. Group differences were assessed using a Mann-Whitney test. *** *p* <0.0001.

Next, we performed a ROC curve analysis based on the binary and discriminating nature of the OPA result, to determine the PIC (Figure 3). For each ROC curves, the area under the curve (AUC) shows at least good analytical performance (AUC>0.70) or even excellent ability to discriminate a positive response by OPA (AUC>0.80).

**Figure 3.**
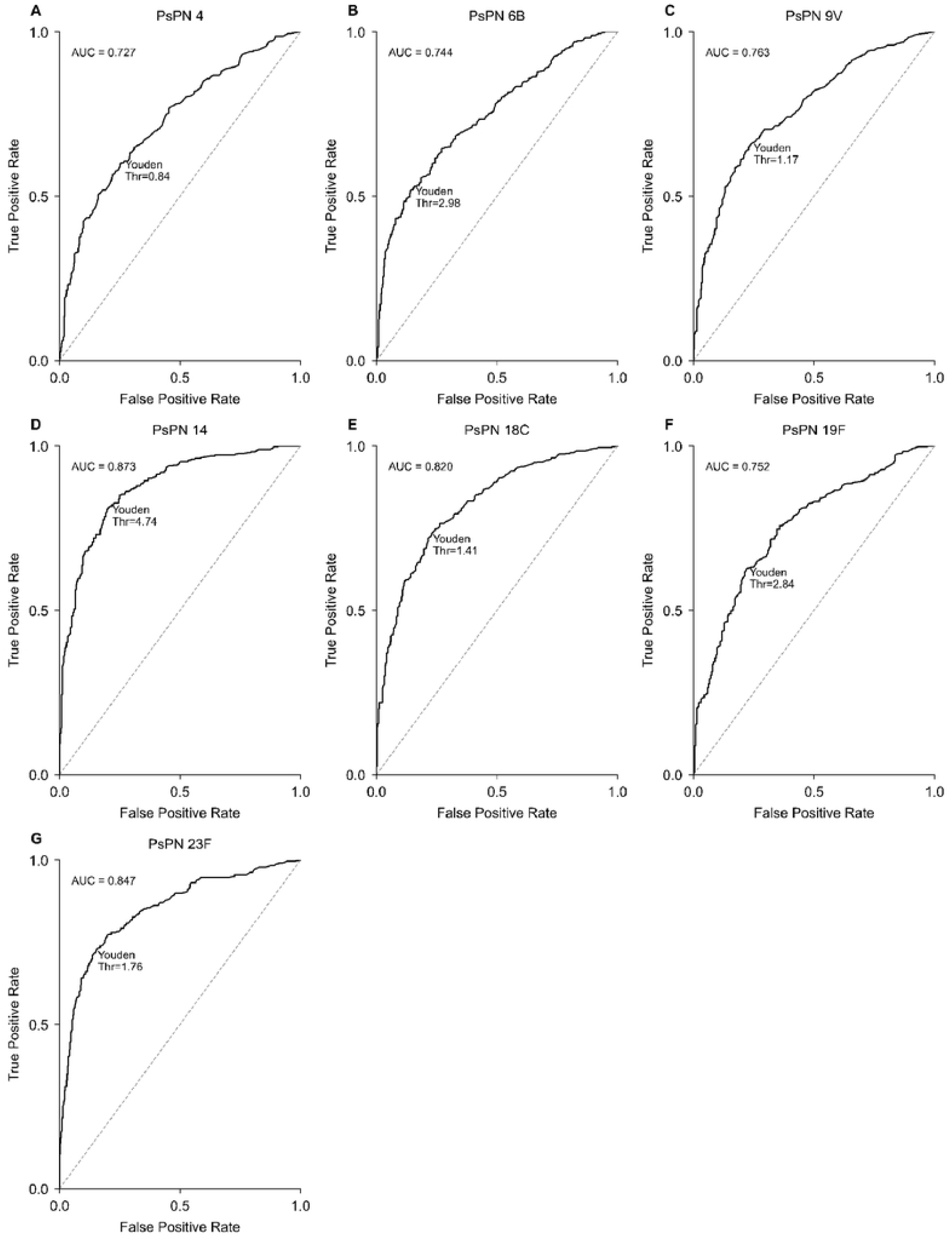
ROC analysis of ELISA lgG concentrations predictive (PIC) of OPA positivity. Receiver operating characteristic (ROC) curves evaluating the performance of serotype-specific ELISA lgG concentrations predictive of OPA positivity (PIC). Optimal thresholds were determined using the Youden index. Panels A-G correspond to serotypes PsPN4, 6B, 9V, 14, 18C, 19F, and 23F.

In order to obtain the most accurate estimate of the PIC, the Youden-index method was applied. The concentration calculation method using Youden’s J index (J=sensitivity + specificity−1) for each serotype offers the best compromise between sensitivity and specificity (Table 3). PIC varied between serotypes, ranging from 0.84 μg/mL for serotype 4 to 4.74 μg/mL for serotype 14, confirming serotype heterogeneity previously observed.

**Table 3.**
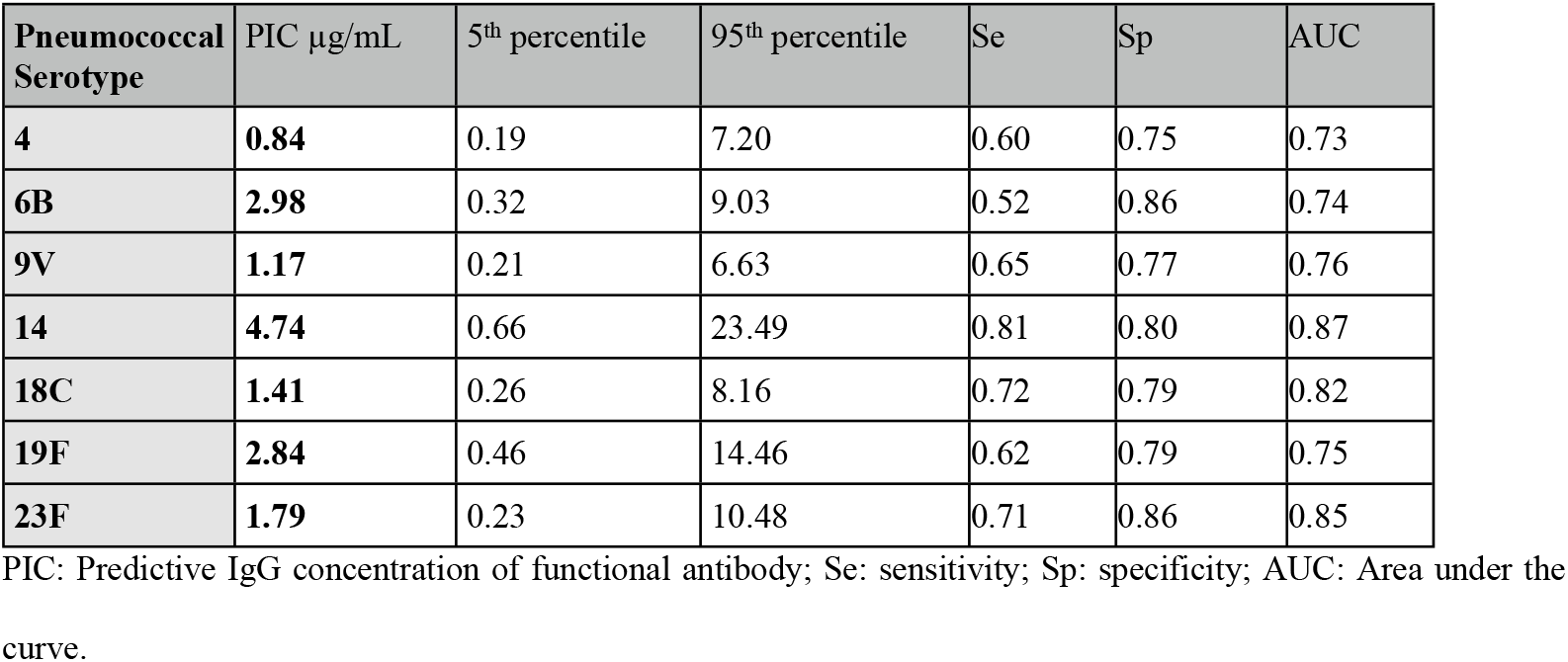
Predictive anti-polysaccharide IgG concentrations determined from ROC curves and Youden-index.

### Bootstrap analysis and evaluation of heterogeneity

To statistically quantify the heterogeneity of PICs between serotypes, it is essential to work on a median PIC rather than a single value (determined by ROC curve analysis). We applied a bioinformatics analysis based on Bootstrap, which simulate the repetition of an experiment by resampling with replacement. Using data from the cohort of 729 samples, we performed the analysis of data from 1,000 resamples, which shows the best compromise for obtaining sufficient statistical power. For each draw, we reproduced the ROC curve analyses and determined the median PIC for each serotype using the Youden index method (Table 4).

**Table 4.** Median PIC determined after 1,000 bootstrap resampling.

| Serotype | Median PIC (ug/mL) |  |
| --- | --- | --- |
|  | Median | 95% CI |
| 4 | 0.88 | 0.51-1.32 |
| 6B | 2.84 | 1.85-3.47 |
| 9V | 1.20 | 0.99-1.74 |
| 14 | 4.71 | 4.08-6.70 |
| 18C | 1.38 | 1.26-2.13 |
| 19F | 2.78 | 1.87-3.08 |
| 23F | 1.68 | 1.42-2.47 |

PIC: Predictive IgG Concentration; CI; confidence intervals (5<sup>th</sup> percentile-95<sup>th</sup> percentile)

Median PIC varied between serotypes, ranging from 0.88 μg/mL for serotype 4 to 4.71 μg/mL for serotype 14, and were very close to values obtained before bootstrap. Serotype heterogeneity of median PIC was confirmed by Kruskal-Wallis’s test indicating that PIC for each serotype are statistically different to each other (H(6)=6398, p<0.0001).

Pairwise comparisons of PIC validated the heterogeneity of immune response depending on the serotype exposed (Figure 4). Comparison between PIC of serotypes 6B and 9V demonstrated marked heterogeneity (p=0.0004). This heterogeneity was even more significant for the serotypes from PCV-7 when data were compared in pairs (p<0.0001).

**Figure 4.**
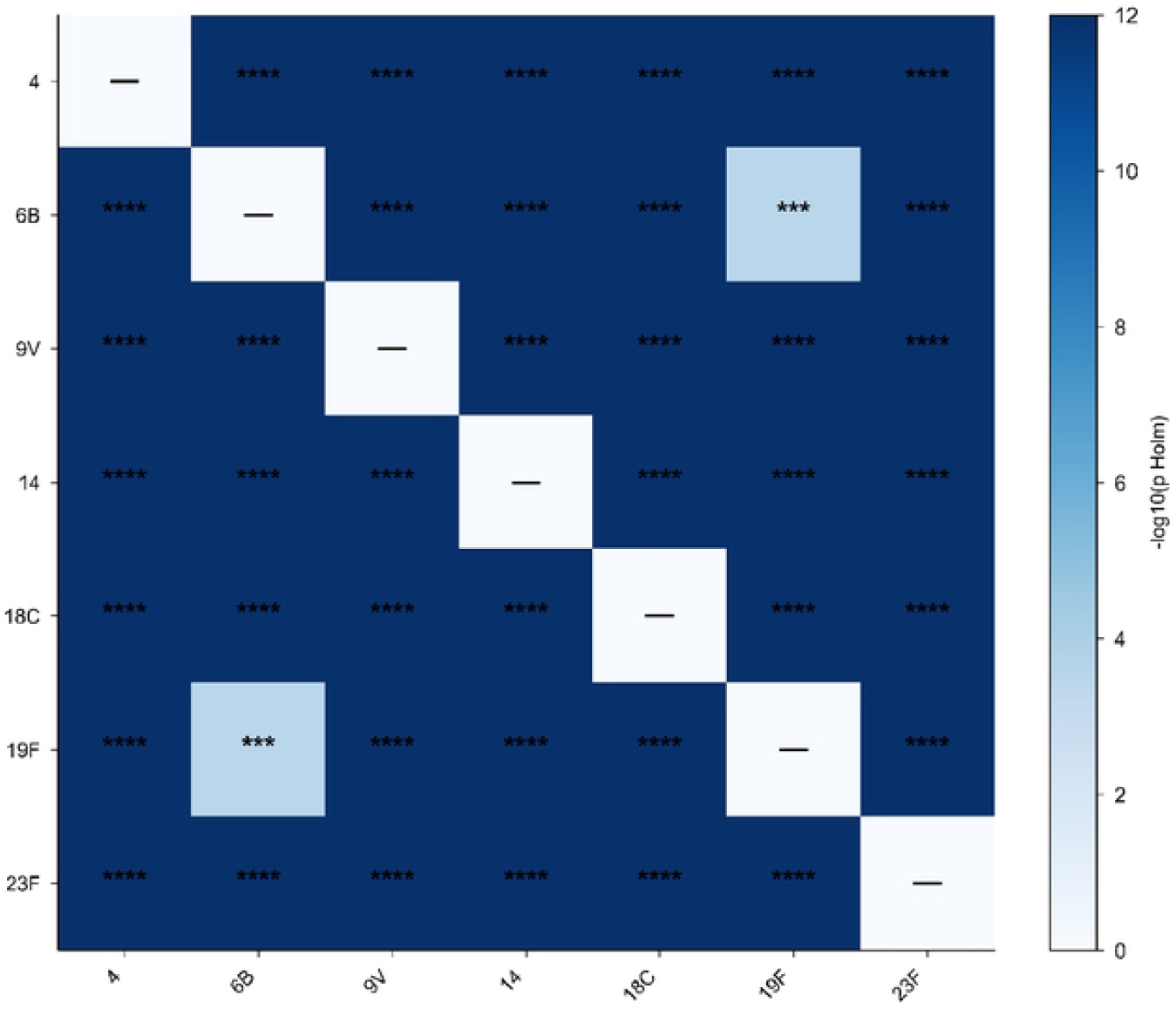
Heterogeneity in serotype-specific PIC threshold determination across pneumococcal serotypes. Heatmap showing pairwise heterogeneity in serotype-specific PIC thresholds estimated from ROC analyses. Variability was assessed using a bootstrap resampling approach (1,000 iterations), in which Youden indices were recalculated across resampled datasets to evaluate the stability of optimal threshold determination. Color intensity reflects the magnitude of heterogeneity between serotype pairs. Statistical significance is indicated in the matrix(*** p<0.001; **** p<0.0001).

### Vaccine status based on thresholds used

The variable distribution of IgG concentrations between serotypes (already observed by Park et al (26)) has not been taken into account by the WHO or AAAAI guidelines recommending a single threshold of 1 or 1.3 µg/mL, respectively. This prompted us to know if the heterogeneous distribution of IgG concentration according to the serotype could have an impact on the conclusions about the vaccine protection. We compared patient vaccination status using the WHO and AAAAI single thresholds with the serotype specific PIC and the 95th percentile of PIC obtained by using the bootstrap method while maintaining the criterion of 70% serotype responders (Table 5).

**Table 5:** Diagnostic of protection according to the threshold of IgG concentration.

| **IgG concentrations** | **PIC** | **95<sup>th</sup>**<br>**percentile**<br>**PIC** | **WHO** | **AAAAI** |
| --- | --- | --- | --- | --- |
| **In favor of protection** | 229 | 160 | 412 | 235 |
| **Not in favor of protection** | 500 | 569 | 317 | 404 |

Patients are in favor of protection when they had at least 70% of responders' serotypes; i.e. 0 to 4 responders serotype for the group 'not in favor of protection' and 5 to 7 responders' serotypes for the group 'in favor of protection'.
Pairwise comparison using McNemar test with Holm correction was applied and for each comparison it demonstrated a *p-value* < 0.0001.

For patients with 0 to 4 responder serotypes, results were not in favor of a protection, whereas for patients with 5 to 7 responder serotypes, results were in favor of a protection. Applying a common threshold for all serotypes (WHO or AAAAI) yielded a higher number of results in favor of a protection, unlike applying serotype-specific concentrations. The Cochrane test on matched proportions has been showed significant in the number of patients considered “in favor of protective vaccine immunity” according to the threshold applied (Q(3)=500.33, p<0.0001). We then pair-wise compared each protection status using a McNemar test based on the threshold applied and demonstrated highly significant changes in protection status due to this immune heterogeneity (p<0.0001 for each comparison), which was highlighted on the basis of the pneumococcal serotype.

## Discussion

By analyzing specific IgG concentrations based on the binary result distribution of the OPA, we were able to determine a predictive IgG concentration (PIC) for each serotype in terms of the functionality of the antibodies investigated in serum samples. This concentration, determined using ROC curves, represented a compromise between sensitivity and specificity in terms of the probability of an antibody titer being effective *in vivo*. The AUC values ranged from 0.73 to 0.87, indicating satisfactory diagnostic performance. Analysis of the distribution of specific IgG concentrations between patients with positive OPA and patients with negative OPA highlighted heterogeneity in the immune response depending on the vaccine serotype studied. This heterogeneity was statistically confirmed *via* dispersion analyses of PICs determined by the dispersion of Youden indices calculated using the bootstrap method. The heterogeneity of the response and protection provided by pneumococcal vaccines can be explained by various factors. For example, the serotype 14 was known for its high immunogenicity, due to its polysaccharide structure. The capsule polysaccharides are composed of repeating units of galactose and N-acetylglucosamine, which are motifs that are very recognized by the immune system(27). The capsule is not particularly thick or mucoid, making the epitopes accessible for effective opsonization. The serotype 14 polysaccharide thus effectively stimulates helper T cells, resulting in high production of specific IgG and robust immune memory(27).

Although enrolling 729 patients, the study only included adult patients. PIC determined for adults will not necessary be similar as that found in pediatric population, which is particularly at risk for IPDs and is one of the most important targets of vaccination campaigns. The high degree of heterogeneity demonstrated in antibody titers was even more pronounced when the immune response against the serotype was effective, as observed by a positive OPA test.

The PICs we have defined did not constitute thresholds of positivity that can be clinically used. They were simply a concentration predicting antibody functionality in a cohort of patients with highly variable clinical characteristics at a given time. PICs differed from the serotype-specific thresholds established by *Park et al* (26), which corresponded to vaccine response thresholds, to track the dynamics of the response before and after vaccination in healthy individuals.

In the daily clinical practice, we proposed an analysis of the seven PCV-7 serotypes and applied the WHO thresholds, *i*.*e*., a diagnosis in favor of a vaccine protection in cases of IgG concentrations of at least five serotypes above 1 µg/mL. However, for some patients with 3 or 4 responder serotypes, considering serotype heterogeneity could be an advantage in refining the diagnosis for these patients, and the subsequent need to develop OPA analyses.

## Conclusion

Determining PICs using Youden indices and the bootstrap method highlighted the heterogeneity of the immune response against pneumococcus vaccine serotypes. The calculation of AUC with values ranging from 0.73 to 0.87 demonstrated high diagnostic performance, leading us to consider the AUCs as potential thresholds for evaluating the pneumococcus vaccine response. Although more complicated to perform technically than ELISA, the OPA test remained indispensable and could not be replaced, highlighting the complementarity of both tests. The heterogeneity of the immune response depending on the serotype raises questions about the relevance of a single threshold. Rather than determining a single threshold, taking into account the heterogeneity of the immune response depending on the serotype could be an alternative issue for better characterization of the anti-pneumococcal vaccine response.

## Materials and methods

### Samples

Between January 2018 and June 2024, 729 samples from adult patients at risk of IPD were investigated in both ELISA and OPA for the 7 serotypes of PCV-7 (4, 6B, 9V, 14, 18C, 19F and 23F). These explorations were intended to determine whether a patient developed sufficient vaccine protection at a given time. The requirements for a protection from IPD were met for vaccine response against more than 70% of the serotypes tested, in line with the WHO’s thresholds.

### Enzyme-linked immune-adsorbent assay

Polysaccharide-specific IgG antibodies titers were measured by ELISA following analytical WHO-recommendations (www.vaccine.uab.edu) as previously described(28). Briefly, pneumococcus polysaccharides (SSI Diagnostica, Hillerød, Denmark) were coated in 96-wells plates (Greiner Bio-One^**®**^, Les Ulis, France). Samples, controls and standard 007-sp (University of Alabama, AL, USA) were pre-absorbed with 5 μg/mL pneumococcal Cell Wall polysaccharide (SSI Diagnostica) and with 10 μg/mL serotype 22F polysaccharide (SSI Diagnostica), then incubated 2 hrs in coated plates. After washing, plates were incubated with alkaline phosphatase-conjugated anti-human goat IgG antibody (SouthernBiotech, AL, USA) for 2 hrs at room temperature. IgG antibodies were detected with *p*-nitrophenyl phosphate substrate (Euromedex, Souffelweyersheim, France). The absorbance was read at 405 nm by a Multiskan FC^**®**^ spectrophotometer (ThermoFisher Scientific, Asnières-sur-Seine, France). Anti-pneumococcus antibody titers were determined in each specimen by analysis of linear regression plots compared with the reference sample 007sp(29).

### Opsonophagocytic assay

OPA was performed for seven specific serotypes (4, 6B, 9V, 14, 18C, 19F, and 23F) to determine functional antibody responses and measured according to the WHO-recommended method (www.vaccine.uab.edu), as previously described(28). Briefly, all serum samples were heat-decomplemented and submitted for their spontaneous bactericidal effect prior to OPA. In case of bactericidal effect, the sample was discarded. Serum samples were serially diluted in round-bottom 96-well plates (Corning, Boulogne-Billancourt, France) and incubated with bacteria (∼50,000 colonies forming unit (CFU)/mL) for 30 min at room temperature. Complement (GentaurFrance, Paris, France) and HL60 cells (ATCC, VA, USA) that had been differentiated to phagocytes (0.5 million) were added to each well. After a 45-min incubation (37°C, 5% CO_2_, 700 rpm), the plates were incubated on ice for 20 min. Bacteria of each well were spotted onto Todd-Hewitt broth-yeast extract agar plates. After an overnight incubation at 37°C, the number of CFU in the plates was enumerated using NICE^**®**^ software (ftp://ftp.nist.gov/pub/physics/mlclarke/NICE/). Opsonization indices (OI) were calculated as interpolated reciprocal serum dilution that killed 50% of bacteria in the assay. The assay sensitivity is the lowest dilution of sera tested (limit of detection: LOD), which is usually 4 for undiluted sera, and the same for each serotype. However, to more precisely quantify functional antibodies, validation of the assay was determined for each serotype-specific assay by low limit of quantification (LLOQ). OPA positivity indexes were determined by each laboratory.

### ROC curve and Youden index

For each pneumococcus serotype, in order to determine the optimal concentration of specific IgG measured by ELISA predictive of a positive OPA response, which will be referred to using the abbreviation PIC, we used the ROC curve method. The samples were categorized according to the OPA result: Positive if greater than or equal to the LLOQ and Negative if less than the LLOQ. For each sample, we calculated sensitivity and specificity, then plotted the ROC curve (sensitivity on the y-axis, 1–specificity on the x-axis). PIC can be determined using the Youden J index (J=sensitivity + specificity−1). It is calculated for each sample, the highest index corresponding to the PIC. The coefficient of variation of the concentration was also calculated.

### Bootstrap

ROC curves and the Youden index can be used to determine a PIC by serotype, that cannot make possible comparison tests between serotypes. To overcome this limitation, we used the bootstrap method. This approach involves generating new samples by randomly drawing samples from the cohort with replacement, then recalculating the PIC at each time. Repeated a defined number of times per serotype, this process yields a PIC distribution that reflects the variability of the estimate. For each serotype, we applied bootstrapping with replacement, performing 10, 100, 1000, and 10,000 resamplings of the same size as the initial cohort. We chose the threshold of 1,000 resampling as the best compromise between statistical power and biological relevance for analyzing PIC heterogeneity for each serotype.

### Normality test

Shapiro-Wilk test did not validate the normality of the distribution of specific Ig concentration values by ELISA, which led us to use nonparametric tests.

### Kruskal-Wallis and Dunn’s post-hoc test with Holm correction

PIC determined by bootstrap were non-normally distributed for all serotypes (Shapiro-Wilk, *p*<0.0001), non-parametric methods were performed for group comparisons. Overall heterogeneity across serotypes was assessed according to Kruskal-Wallis test and, in cases of significant results, we applied a Dunn’s post-hoc test with Holm correction to identify pairwise differences for multiple testing. Holm-adjusted *p* values <0.05 were considered statistically significant.

### Statistical analysis

All statistical tests were performed using Python^®^ software V3.10 (Wilmington, DE, United States) with the following plugins: Pandas^®^ V1.5.3, Sklearn, matplotlib^®^ 3.6.3, Scipy^®^ 1.10.1, Scikit-learn 1.2.1^®^, Scikit-posthocs 0.11.4^®^, Statsmodels 0.14.4^®^ and Numpy^®^ 1.24.1. The applied test was introduced in the legends of the figures and tables.

## Notes

**Funding:** This work was supported by grants from AP-HP, GHU Paris Centre, University Paris Cité and INSERM.

**Competing interests:** The authors state no conflict of interest.

### Competing Interest Statement

The authors have declared no competing interest.

